# BAMBI: Integrative *b*iostatistical and *a*rtificial-intelligence *m*odels discover coding and non-coding RNA genes as *bi*omarkers

**DOI:** 10.1101/2024.01.12.575460

**Authors:** Peng Zhou, Zixiu Li, Feifan Liu, Euijin Kwon, Tien-Chan Hsieh, Shangyuan Ye, Shobha Vasudevan, JungAe Lee, Chan Zhou

## Abstract

Accurate disease diagnosis and prognosis are crucial for effective treatment management and improving patient outcomes. However, accurately detecting early signs of certain diseases or recurrence remains challenging. Existing machine-learning methods for identifying gene expression biomarkers have several limitations, including poor performance on independent test datasets, inability to directly process omics data, and difficulty in identifying noncoding RNA genes as biomarkers. Additionally, these methods may not provide sufficient biological interpretation of their results, and the panel biomarkers they identify may not be suitable for clinical application. To address these limitations, we have developed a new computational method called BAMBI, which integrates multiple machine-learning algorithms and statistical approaches to identify putative coding and noncoding genes as biomarkers for disease diagnosis and prognosis. We evaluated BAMBI ability to identify diagnostic and prognostic biomarkers by analyzing multiple RNA-seq datasets from cancerous and non-cancerous diseases at population levels. The results from BAMBI demonstrate significant biological interpretability and state-of-the-art prediction performance. When the singular gene identified by BAMBI is used as a diagnostic biomarker, it achieves a balance accuracy exceeding 95% in studies of both breast cancer and psoriasis. Additionally, the prognostic biomarkers that BAMBI identifies from RNA-seq data of Acute Myeloid Leukemia (AML) patients significantly correlate with the survival rates in an independent AML patient cohort. Additionally, BAMBI outperforms existing methods by delivering more robust results, identifying biomarkers with fewer genes, and simultaneously achieving superior prediction accuracy. We have implemented BAMBI into user-friendly software for the research community. In summary, BAMBI serves as a more reliable pipeline for identifying both coding and noncoding genes as biosignature markers, enhancing the accuracy of disease diagnosis and prognosis. BAMBI is available via https://github.com/CZhouLab/BAMBI.

## INTRODUCTION

Molecular biomarkers have been widely used to assist in diagnosis and prognosis. There are four major types of molecular biomarkers at the biological level: DNA, RNA, protein, and metabolic biomarkers (FDA, 2016). RNA biomarkers have several advantages over other types of molecular biomarker: 1) The rapid and drastic changeability of RNA expression provides real-time monitoring of disease status and treatment response (Byron et al., 2016; FDA, 2016); 2) the RNA biomarker may provide insights into active gene expression and their roles in disease manifestation and progression(Xi et al., 2017a); 3) RNA molecules shows high tissue of origin specificity compared to DNA biomarkers, such that RNA biomarkers becomes especially useful in cancer diagnosis and prognosis (Xi et al., 2017b); 4) multiple types of RNA (including mRNAs and lncRNAs) can be detected in bodily fluids, such that RNA biomarkers can be applied in the non-invasive samples which will increase patient comfort and compliance; 5). The high specificity of RNA biomarkers contributes to the advancement of personalized medicine by optimizing clinical trial processes and accelerating the approval of new drugs. Additionally, advances in next-generation RNA sequencing (RNA-seq) techniques make RNA biomarker identification and quantification more reliable compared to the traditional microarray techniques.

However, the existing methods for identifying RNA biomarkers have several limitations. 1) The biomarkers identified by them have poor predictive performance when applied to independent test datasets. This issue not only undermines confidence in these biomarkers but also raises concerns about their clinical applicability. 2) they are unable to directly process RNA-seq data, limiting their utility for genome-wide quantification of RNA molecules. 3) they do not consider non-coding as potential biomarkers, despite evidence that non-coding RNAs may exhibit a higher degree of disease-specific expression compared to mRNAs (Ratti et al., 2020). This could lead to missed opportunities for novel diagnostics and treatments that non-coding RNA biomarkers might present. 4) Existing methods may not provide sufficient insights into the biological relevance behind their findings, limiting understanding of the underlying disease mechanisms. 5) While biomarkers encompassing more genes generally lead to improved performance, existing methods falter by either delivering poor performance or incorporating an excessively large number of genes. This over-inclusion, often a result of overfitting in training data, renders the biomarkers impractical for clinical use due to increased costs, extended time for results, and added complexity in analysis and interpretation. Furthermore, a test panel with an abundance of biomarker genes necessitates larger sample volumes, which may not be feasible in situations with limited sample availability. Additionally, a higher number of biomarker genes increases the risk of false readings and complicates the validation process. Therefore, the challenge remains to identify an optimal number of RNA biomarker genes while still reserving high prediction performance.

To overcome these challenges and limitations, we developed BAMBI (*<u>B</u>*iostatistics and *<u>A</u>*rtificial-Intelligence integrated *<u>M</u>*ethod for *<u>B</u>*iomarker *I*dentification), a robust pipeline that identifies both coding and non-coding RNA biomarkers for disease diagnosis and prognosis. BAMBI can process RNA-seq data and microarray data to pinpoint a minimal yet highly predictive set of RNA biomarkers, thus facilitating their clinical application. Additionally, BAMBI offers visualization of biomarker expression and interpretation their functions using co-expression networks and literature mining, enhancing the interpretability of the results.

## 2 MATERIALS AND METHODS

### 2.1 Data source

#### RNA-seq data

We collected two RNA-seq data sets for this study: 1) the TCGA breast cancer dataset and 2) the psoriasis dataset. In the TCGA breast cancer dataset, we downloaded the transcriptomics data in BAM format of 116 solid ductal and lobular neoplasms biospecimens and 112 adjacent normal solid tissue samples from the TCGA-BRCA project (Weinstein et al., 2013). All these biospecimen were collected from 112 breast cancer patients. Because a few patients provide more than one tumor biospecimens, it results in a few more data of tumor biospecimens than the normal sample. We converted the downloaded BAM data into Fastq format using the BIOBAMBAM2 software (Tischler & Leonard, 2014). The RNA-seq data of psoriasis dataset was downloaded from GEO database (accession number: GSE54456). These data were obtained in SRA format and then we used the fastq-dump toolkit from NCBI website to convert SRA file into Fastq format.

#### Microarray data

Two microarray datasets were used in this study. The first one is the Colon Cancer dataset from (Alon et al., 1999), which was originally utilized to differentiate between cancerous and non-cancerous colon tissues, comprising data from 62 patients (40 with tumors and 22 with normal tissues). The second one, Prostate Cancer dataset, comes from (Singh et al., 2002). This dataset was initially used to examine the differences in gene expression between tumor and non-tumor prostate samples, including information from 102 patients (52 with tumors and 50 normal). For compatibility with the BAMBI software, we transformed the microarray data into the required input format. The processed tables are accessible on our GitHub repository.

### 2.2 Gene annotation

We used LncBook Version 2.0 (Ma et al., 2019) as the mapping reference genome annotations for RNA-seq data. It contains the genomic location annotation of 19957 coding genes and 101293 lncRNA genes.

### 2.3 Feature selection embedded in BAMBI

Sequencing data are normally high-dimensional data with a large number of features but a small number of samples. In machine learning, high-dimensional data are prone to overfitting because the abundance of features relative to few numbers of observations can lead models to detect spurious patterns that do not generalize well to unseen data. In BAMBI, we applied a two-step feature selection, that are statistical based feature selection and Machine learning based feature selection, to decrease the data dimension.

#### 2.3.1 Statistical based feature selection

We employed a suite of statistical methods tailored to the pragmatic demands of biomarker identification. These include differential expression analysis, fold change analysis, extremely lowly expressed genes filter and high distribution overlap genes filter as follows:

##### (1) Differential expression analysis

We adopted the Wilcoxon-Mann-Whitney test as the default method for RNA-seq differential expression analysis in BAMBI. This is because Wilcoxon rank-sum test, a non-parametric method has been shown to accurately differentially expressed (DE) genes in RNA-seq data of samples larger than 8 (Li et al., 2022). Compared to the conventionally DE analysis methods (DESeq2 (Love et al., 2014) and EdgeR (Robinson et al., 2009)), Wilcoxon-Mann-Whitney test shows its robustness to outliers and consistent control of false positive rates under desired thresholds for population level of studies (Li et al., 2022). Whereas we used the limma method (Ritchie et al., 2015), to perform differential expression analysis for microarray data. Limma has been integrated into the BAMBI pipeline to do differential expression for microarray data.

##### (2) Fold Change analysis

Fold change represents the ratio of expression levels or quantities between two conditions, often used to measure changes in gene expression in biological experiments (Goss Tusher et al., n.d.). Its significance in biomarker detection lies in its ability to discern notable alterations, providing a straightforward and intuitive measure of effect size. In BAMBI, we calculate the fold change score based on the absolute log2 of the gene median expression levels ratio between different conditions.

##### (3) Extremely lowly expressed genes filtering

Removing extremely low expression genes is a critical step in biomarker detection. Genes with consistently extremely low expression across samples often contribute to noise rather than signal in the data. We excluded the genes which maximum expression levels across all samples is less than a certain threshold. Because different types of gene have different expression scale. Non-coding RNA genes are commonly expressed much lower than coding genes, so we set FPKM=1.0 as the threshold for coding gene and set FPKM =0.01 as the threshold for lncRNA genes.

##### (4) High distribution overlap genes filtering

Unlike traditional methods that rely on mean or median differences to evaluate gene expression heterogeneity, BAMBI introduces an additional criterion by estimating the distribution overlap area between groups. This approach offers a more comprehensive and accurate representation of gene expression heterogeneity between groups. The expression distribution was estimated using the kernel density estimate, implemented via the Python package *KDEpy*. Then the size of overlap area between expression distributions of two groups (e.g., patients and controls) was calculated using the python package *numpy*.

#### 2.3.2 Machine learning based feature selection

Accurate prediction is a key attribute of an effective biomarker. (Rifai et al., 2006). In the statistical based feature selection stage, we identify candidate genes through group-level statistical differences. However, this step alone might not assure precise disease prediction for individual samples. To enhance this, BAMBI also incorporates a machine learning based feature selection, focusing on enhancing prediction accuracy at the individual sample level. We integrated the following strategies into the BAMBI tool for the feature selection when building the machine learning models:

##### (1) Recursive Feature Elimination: Optimizing Gene Selection

We employ the recursive feature elimination (RFE) for feature selection to identify the optimal gene set. RFE stands as a widely recognized algorithm for this purpose. Initially, the model is trained on the entire gene set, and the significance of each gene for prediction is determined. Subsequently, genes deemed least important are excluded, while concurrently recording the model’s prediction performance based on the remaining genes. This pruning process is repeated iteratively until only a single gene remains. By plotting these recorded performances against gene count, a curve is generated. The optimal gene set is determined by identifying the knee point of this curve. We have selected the SHAP algorithm (Lundberg et al., 2017) to evaluate the genes predictive significance for RFE. Renowned for its game theory foundation and lucid interpretation, SHAP can be integrated with almost all conventional machine learning and deep learning models. In contrast, another cutting-edge machine learning interpretation method, LIME, can sometimes exhibit instability. Its performance is contingent upon specific parameter settings, such as the neighborhood kernel width.

##### (2) Enhanced Ten-Fold Cross-Validation for Robust Feature Selection

We built a structure similar to the ten-fold cross-validation, a classic machine-learning resampling method, do the feature selection. The entire dataset is separated into ten portions. Because the transcriptomics datasets are commonly unbalanced between different groups of biospecimens, we kept each portion containing approximately the same percentage of samples of each target class as the original dataset. We each time took nine of them as the training set for gene selection and model training and the remaining portion as the testing set. As the result, we generated ten train-test set pairs. Then, we did the feature selection and model training on each training set and evaluated their performance on the relevant testing set.

Conventional approaches employ ten-fold cross-validation (10-CV) for feature selection by assessing and comparing the aggregated 10-CV performance metrics of various feature selection hyperparameters. The hyperparameter that demonstrates the highest aggregated performance is then selected to guide feature selection on the entire training set, ultimately determining the final feature set. However, it may overlook valuable insights derived from the 10-CV process. For instance, a set of features consistently selected across different training and testing pairs could indicate higher robustness and predictive power. Similarly, features frequently chosen by diverse model types which are based on varying assumptions might exhibit greater heterogeneity.

In contrast, our novel method leverages the comprehensive information available from the 10-CV process. We employ a two layers statistical analysis to enhance the prediction of singular RNA biomarkers and a panel of multiple RNAs biomarkers. This approach ensures a more nuanced and thorough utilization of the information obtained from 10-CV.

##### (3) Two Layer Statistical Downstream Analysis of 10-CV

By applying *M* types of models to ten paired training-testing sets of 10-CV process, we generated 10 x *M* trained candidate predictive models. Each model, through its independent feature selection process, potentially selects different gene combinations as predictors. BAMBI collects information from these 10 x *M* candidate models, including the predictor gene information, model type and their relative test set performance.

Based on this information, BAMBI conducts two layers’ statistics to predict singular RNA biomarker and a panel of multiple RNAs biomarker. When predicting the singular biomarker, BAMBI method hinges on a triad of statistical criteria: the number of training-testing set pairs the gene be selected, diversity of model types including the gene, and frequency of gene presence across models. Of these, the ‘covered training-testing set pairs number’ is paramount, as it validates the gene’s performance across diverse data splits, avoiding overfitting to specific sets. A gene’s robustness is further supported if it is consistently selected by diverse models (‘model type number’), suggesting greater group heterogeneity. ‘Frequency’ also plays a role, reinforcing the gene’s discriminative power if it is selected across numerous models.

When predicting a panel of multiple RNAs biomarkers, candidate models are clustered based on identical gene sets and types, with an emphasis on the ‘covered training-testing set pairs number’ and the model’s average test set performance metrics. A model is considered significantly reliable if selected in 70% or more of the training-testing pairs. This priority ranking (‘covered pairs number’ > ‘average test set performance’ > ‘model gene count’) is adjusted when a model’s selection frequency is low (e.g., ‘covered pairs number’ of one to two), shifting importance to ‘average test set performance’. This flexible approach allows for the discernment of the most robust predictive models, accommodating both generalizability and performance.

##### (4) Model Selection and Hyperparameter Optimization in BAMBI

In BAMBI, we have selected a range of machine learning classification models as potential candidates, including support vector machine (SVM), K-nearest neighbor (KNN), logistic regression, and Bayes. These classifiers are prevalent in bioinformatics analyses and have consistently demonstrated commendable performance (Hasan, Alam, et al., 2021; Hasan, Basith, et al., 2021; Hasan et al., 2020).

To select the best model training hyperparameters, we applied the grid search method in BAMBI. For each model, we define an appropriate set of possible hyperparameter combinations. The grid search method exhaustively generates candidates from all possible hyperparameter combinations and evaluates the model based on the ten-fold cross-validation. The hyperparameters combination with the best cross-validation score is used to train the final model.

### 2.4 Generation of Heatmap

We used for R package “pheatmap” with default setting to generate the heatmap of this study.

### 2.5 Co-expression network analysis

We constructed the co-expression networks for lncRNAs using the mcxarray program in the Markov Clustering (MCL)-edge network analysis tool (Walker, 2011) (http://micans.org/mcl/) with Spearman correlation. In this study, we choose 0.8 as the Spearman correlation cutoff to balance the number of singletons and the median node degree as recommended by the MCL protocol.

### 2.6 Gene Ontology (GO) analysis

We used the DAVID Functional Annotation platform (http://david.abcc.ncifcrf.gov/)(Huang et al., 2009; Jiao et al., 2012) to perform the GO enrichment analysis for the protein-coding genes identified in each co-expression cluster. Only protein-coding genes with an FPKM value greater than 1 in at least one sample from the entire dataset were used as the background for GO enrichment analyses.

### 2.7 Survival analysis

Survival analysis was performed used the python package lifelines. The p-value was calculated by the logrank test.

### 2.8 RNA-seq data sets pre-processing for BioDISCML, ILRI, and ECMARKER tools

Given that these three tools are unable to process raw RNA-seq data directly, we adopted a preprocessing approach consistent with BAMBI’s internal function. First, RNA-seq reads were mapped to the reference genome by HISAT2 tool (Kim et al., 2019). Subsequently, the HTSeq tool (Anders et al., 2015) was used to quantify the mapped alignments. Finally, quantified read counts mapping to each RNA gene was used to calculate the expression levels of RNA genes in FPKM. Prior to applying each tool, we converted the expression levels into the specific input format required by each tool.

### 2.9 Comparison of BAMBI with BioDiscML, ILRI, ECMarker

We compared the performance of BAMBI and current alternative methods (BioDISCML (Leclercq et al., 2019), ILRC (Yu et al., 2021), ECMarker (Jin et al., 2021)) in two RNA-seq datasets (breast cancer dataset and psoriasis dataset) and two microarray datasets (colon cancer dataset (Alon et al., 1999) and prostate cancer dataset (Singh et al., 2002). Because the split of train and test data will strong influence the performance score of the target method, especially for small datasets with less than 1000 samples, we applied a cross validation comparison structure to obtain a convincing comparison result for different methods. For each dataset, we did two times of fivefold stratified cross validation separation with different randomization. It resulted in ten different train-test set pairs, each train set including 80% of samples and each test set including 20% of samples. We applied all methods on these ten different train sets to predict putative biomarkers. Then we evaluated the performance of these identified biomarkers on the corresponding test set. The average performance of the ten train-test set pairs was used to compare the performances of BAMBI and other methods.

## RESULTS

### BAMBI, a computational tool for discovering singular and panel RNA biomarkers

The clinical utility of RNA biomarkers in disease diagnosis and prognosis hinges on the accurate detection of both singular and panel RNA biomarkers. In practical scenarios, RNA biomarker detection involves processing sequencing materials obtained from various sources, including whole blood and organ tissues from both patients and healthy individuals. These samples undergo RNA sequencing, producing comprehensive RNA-seq data. Sequence data is subsequently analyzed using the high-throughput sequencing analysis to identify the candidate RNA biomarkers. (**Fig 1A**).

**Fig 1:**
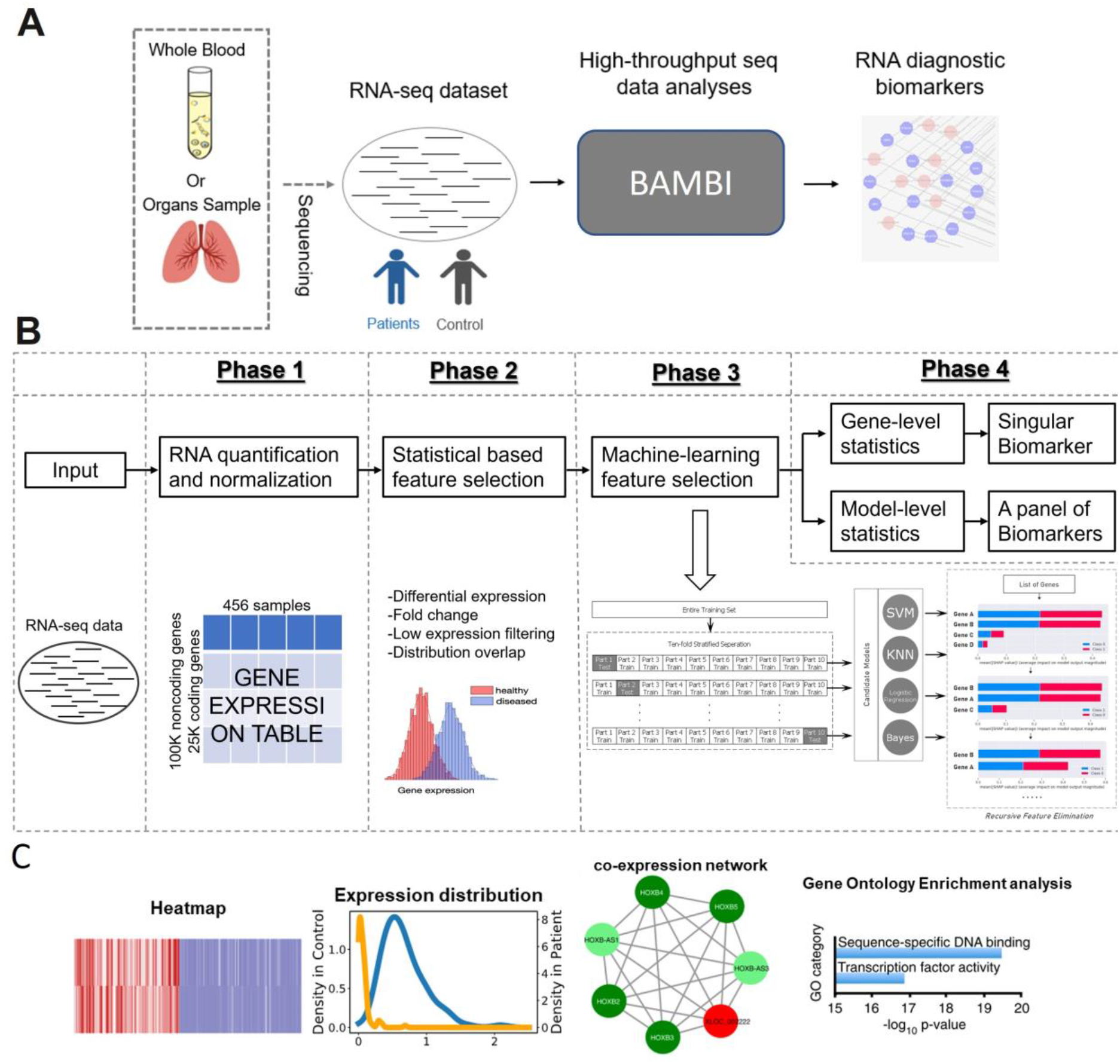
Discovering coding and noncoding RNA biomarkers by integrative biostatistical and artificial-intelligence models. (A) RNA biomarker detection in practical scenarios (B) BAMBI pipeline overview. BAMBI integrates both of statistical methods and machine-learning models. BAMBI can predict both the singular putative biomarker and panel putative biomarkers of coding and noncoding RNA genes. BAMBI integrates biostatistical and machine-learning methods to select expression features. Biostatical approaches significantly reduce the high-dimensions of genes based on expression table. BAMBI combined two major statistical methods: gene differential expression analysis and calculation the overlap area of the expression level distributions among two populations. BAMBI select the coding and noncoding RNA genes as the gene features for the downstream machine-learning models, if the gene is differentially expressed and have a small size of the overlap area of its expression level distributions among two populations. (Pls see Materials and Methods for details). The 10-CV-based recursive feature elimination strategy avoids overfitting and yields more robust selected-features (Pls see Materials and Methods for details). (C) BAMBI automatically discovers and visualizes coding and non-coding RNA biomarkers. Heatmaps and expression level distributions of putative biomarkers are presented to illustrate the expression changes of putative biomarkers between different groups, e.g., patients and health controls. Co-expression network graph along with the Gene Ontology (GO) enrichment bar-plot are also presented for the putative lncRNA biomarkers to indicate their potential functional roles in related disease.

In addition to coding RNA (mRNA) biomarkers, long non-coding RNAs (lncRNAs) have also been implicated as putative biomarkers for disease diagnosis and progression, due to their critical roles in pathogenesis of various diseases and their dysregulated gene expression levels. LncRNA is recognized as promising RNA biomarkers for the development of non-invasive liquid biopsy biomarkers for disease(Beylerli et al., 2022; Jiang et al., 2019; Karimi et al., 2022; Wang et al., 2020). The advances of high-throughput sequencing technique have enabled comprehensive studying quantifying relatively lowly expressed lncRNAs to facilitate robust identification of previously unrecognizable and undetectable biomarkers in patients.

In this study, we introduce BAMBI (**Fig 1B**), a novel automated pipeline developed using biostatistical and machine learning approaches. BAMBI is adept at discerning potential biomarkers from RNA-seq or microarray data, encompassing both coding and non-coding RNAs. This tool is uniquely capable of identifying individual RNA biomarkers with high predictive power and biological relevance. Additionally, it can detect a panel of multiple RNAs biomarkers, ensuring a minimal yet effective set that retains significant predictive accuracy. BAMBI is subdivided into four phases:

- *In phase 1*, BAMBI preprocesses the input transcriptomics data (RNA-seq data or microarray data) as described in the materials and methods.
- *In phase 2*, BAMBI employed a suite of statistical methods tailored to the pragmatic demands of biomarker identification. These included differential expression analysis, fold change analysis, low-expression genes filter and high distribution overlap genes filter. BAMBI excluded any feature which are extremely lowly expressed or have high overlap area between the distributions of two groups, or do not have significantly differentiated expressed (please see **Materials and Method** for details).
- *In phase 3*, BAMBI further reduces the features using machine-learning based feature selection. Particularly, we implemented ten-fold cross-validation across the entire dataset, resulting in ten train test set pairs. Within each training set, we employed Recursive Feature Elimination (RFE) embedded with SHAP value(Lundberg et al., 2017), to determine the optimal gene panel. Each model was then trained using this refined gene set and subsequently evaluated against its respective test set.
- *In phase 4*, BAMBI leverages the comprehensive information available from the 10-CV process, and it conducts two layers’ statistics to enhance the prediction of singular RNA biomarker and a panel of multiple RNAs biomarker. This approach ensures a more nuanced and thorough utilization of the information obtained from 10-CV (please see **Materials and Method** for details).

Additionally, BAMBI provides the expression patterns of the discovered putative biomarkers which can be visualized in both heatmap or distribution or bar-plots. Co-expression network and Gene Ontology (GO) enrichment analysis will also be visualized for lncRNA biomarkers to illustrate the potential biological significance of the discovered lncRNA biomarkers (**Fig 1C**).

### BAMBI identifies diagnostic biomarkers with high prediction power and biological significance in breast cancer

We tested BAMBI in discovering mRNA and lncRNA biomarker for diagnosing breast cancer. BAMBI was applied to analyze the RNA-seq data of 116 biospecimens from primary ductal and lobular neoplasms and RNA-seq data of 112 biospecimens from the adjacent normal solid tissues. BAMBI identified both singular and panel biomarkers from this dataset. Here we focused on presenting the results of singular biomarkers identified by BAMBI.

Two mRNA (TSLP and SPRY2) and five lncRNA molecules (HSALNG0022084, HSALNG0119995, HSALNG0112904, HSALNG0075756, HSALNG0116686) were discovered as the putative singular diagnostic biomarkers for breast cancer. Each of these seven RNA biomarkers could be used to facilitate diagnosing breast cancer with high prediction power up to 98.7% of balanced accuracy (see **Fig 2A**). The expression profiles of these seven RNA biomarkers across breast cancer tumors and normal tissues in heatmap and distribution graph (**Fig 2 B-C**) clearly show the significantly distinct expression patterns of the discovered RNA biomarkers between breast cancer tumors and normal tissues.

**Fig 2:**
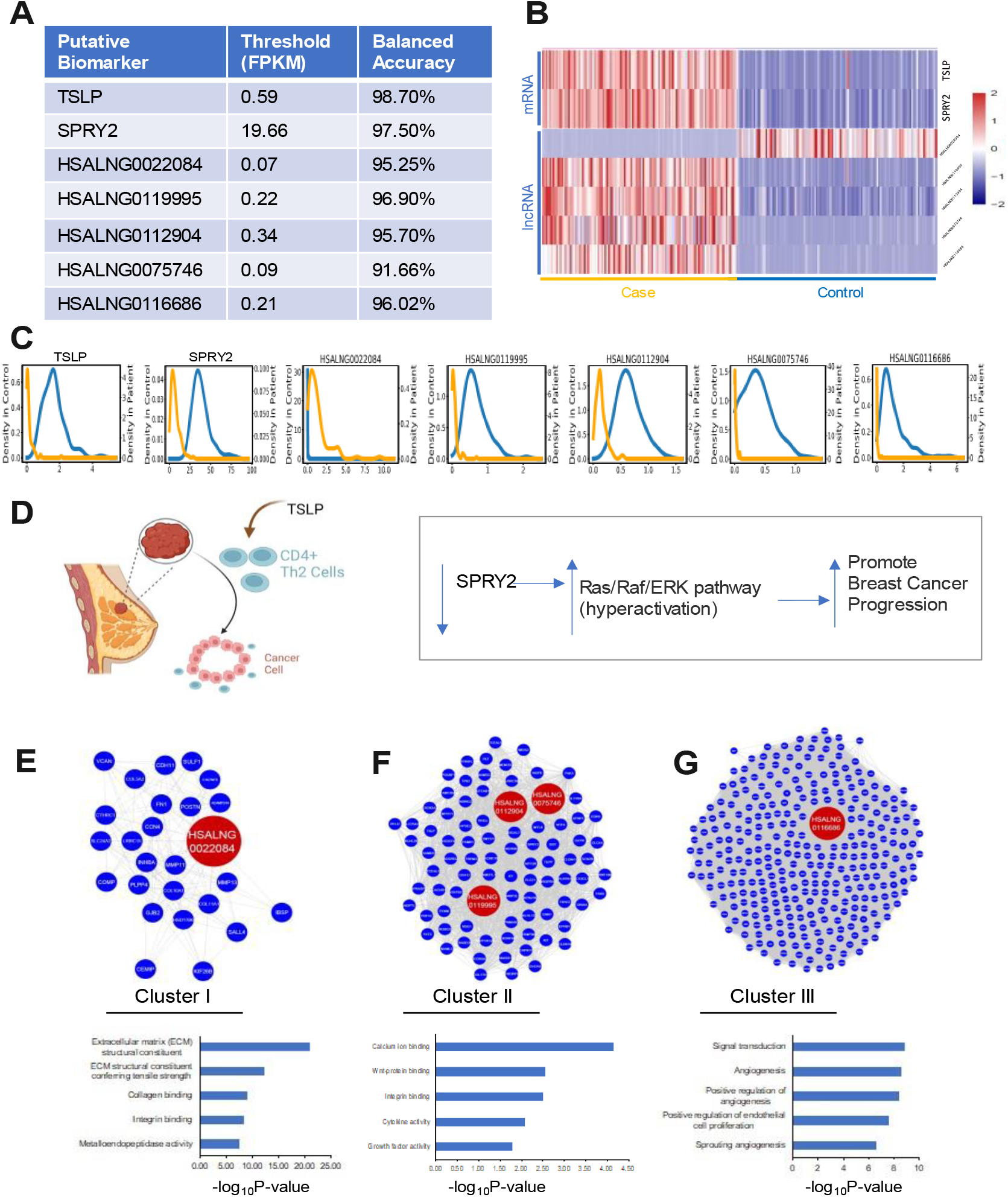
BAMBI discover the biological informative singular mRNA or lncRNA gene as the putative biomarkers for breast cancer diagnosis. We applied BAMBI to the RNA-seq data of the tumor biospecimen and nearby normal biospecimens from the cohort of 112 breast cancer patients. BAMBI found that each of two coding genes (TSLP and SPRY2) and the five lncRNAs (HSALNG0022084, HSALNG0119995, HSALNG0112904, HSALNG0075746, HSALNG0116686) would be predictive for breast cancer diagnosis, each with over 91% balanced accuracy of prediction. (A) The prediction performance for each of the putative biomarker in diagnosing breast cancer. (B) Heatmaps of the expression profiles for these seven candidate putative biomarkers across controls and breast cancer specimens. (C) Expression distributions of for these seven candidate putative biomarkers across controls and breast cancer specimens. (D) Known functional roles of the putative coding gene biomarkers (TSLP and SPRY2) in breast cancer. TSLP were shown to inhibit the breast tumorigenesis through Th2 (Boieri et al., 2022; Demehri et al., 2016b; Guennoun et al., 2022), and the repressing the expression of SPRY2 would lead to promote the breast cancer progression through the hyperactivation of the Ras/Raf/ERK pathway (Faratian et al., 2011; Hanafusa et al., 2002; Kawazoe & Taniguchi, 2019). (E-G) The five putative lncRNA-biomarkers are contained in three co-expression clusters (Cluster I, II and III). The coding genes of the Cluster I, II, III are enriched in the Gene Ontology (GO) categories significantly related to the breast cancer progression. (E) The Cluster I which contains the putative lncRNA biomarker (HSALNG0022084) are enriched in the coding genes involving in the extracellular matrix (ECM) structural constituent. (F) The Cluster II which contains the putative lncRNA biomarker (HSALNG0119995, HSALNG0112904, HSALNG0075746) are enriched in the coding genes involving in the calcium ion binding and Wnt-protein binding. (G) The Cluster III which contains the putative lncRNA biomarker (HSALNG0116686) are enriched in the coding genes involving in the angiogenesis signal transduction.

The seven discovered RNA biomarkers shed light significant biological significance in breast cancer. The TSLP biomarker has been shown to block breast cancer development through the activation of CD4+ T cells (Demehri et al., 2016a) (**Fig 2D, left**). And, the other mRNA biomarker, SPRY2, has also been shown to negatively regulate breast cancer progression. Repressing the expression of SPRY2 will hyperactivate Ras/Raf/ERK pathway, leading to promote breast cancer progression (Figure 2D, right). The five lncRNA biomarkers are co-expressed with three groups of coding-genes (**Fig 2E and F**). Among these five lncRNA biomarkers: 1) the HSALNG002284 was co-expressed with coding genes involved in extracellular matrix structural constituent, collagen binding and integrin binding (**Fig 2E and F, left**). 2) These three lncRNA biomarkers (HSALNG0119995, HSALNG0112904, HSALNG0075756) are co-expressed with coding genes involved in calcium or Wnt-protein binding (**Fig 2E and F, middle**). 3) The other one lncRNA biomarker (HSALNG0116686) is co-expressed with coding genes participating in the angiogenesis signal transduction (**Fig 2E and F, right**). It indicates that the coding genes of these three co-expression clusters (**Fig 2E and F**) with the putative lncRNA biomarkers commonly contribute to the breast cancer progression.

### BAMBI identifies diagnostic biomarkers with high prediction power and biological significance in psoriasis

To evaluate the ability of BAMBI in discovering diagnostic biomarkers in non-cancerous diseases, we applied BAMBI to analyze the RNA-seq data of 174 psoriasis biospecimen from 92 patients and 82 healthy controls. BAMBI discovered both singular and panel biomarkers from this transcriptomics dataset of psoriasis patients and healthy controls. We present the results of panel biomarkers in the section of comparing with other methods, here we focused on presenting the results of singular biomarkers identified by BAMBI.

Two mRNA biomarkers (S100A9 and S100A7) and two lncRNA biomarkers (HSALNG0002446 and HSALNG0129303) were discovered as putative biomarkers. Each of these four RNA biomarkers could be used to diagnose psoriasis with balanced accuracy over 98% (**Fig 3A**). The expression profiles of these four biomarkers (**Fig 3B-C**) in heatmap and distribution plot depict the strictly difference expression patterns of these four biomarkers between psoriasis patients and healthy controls.

**Fig 3.**
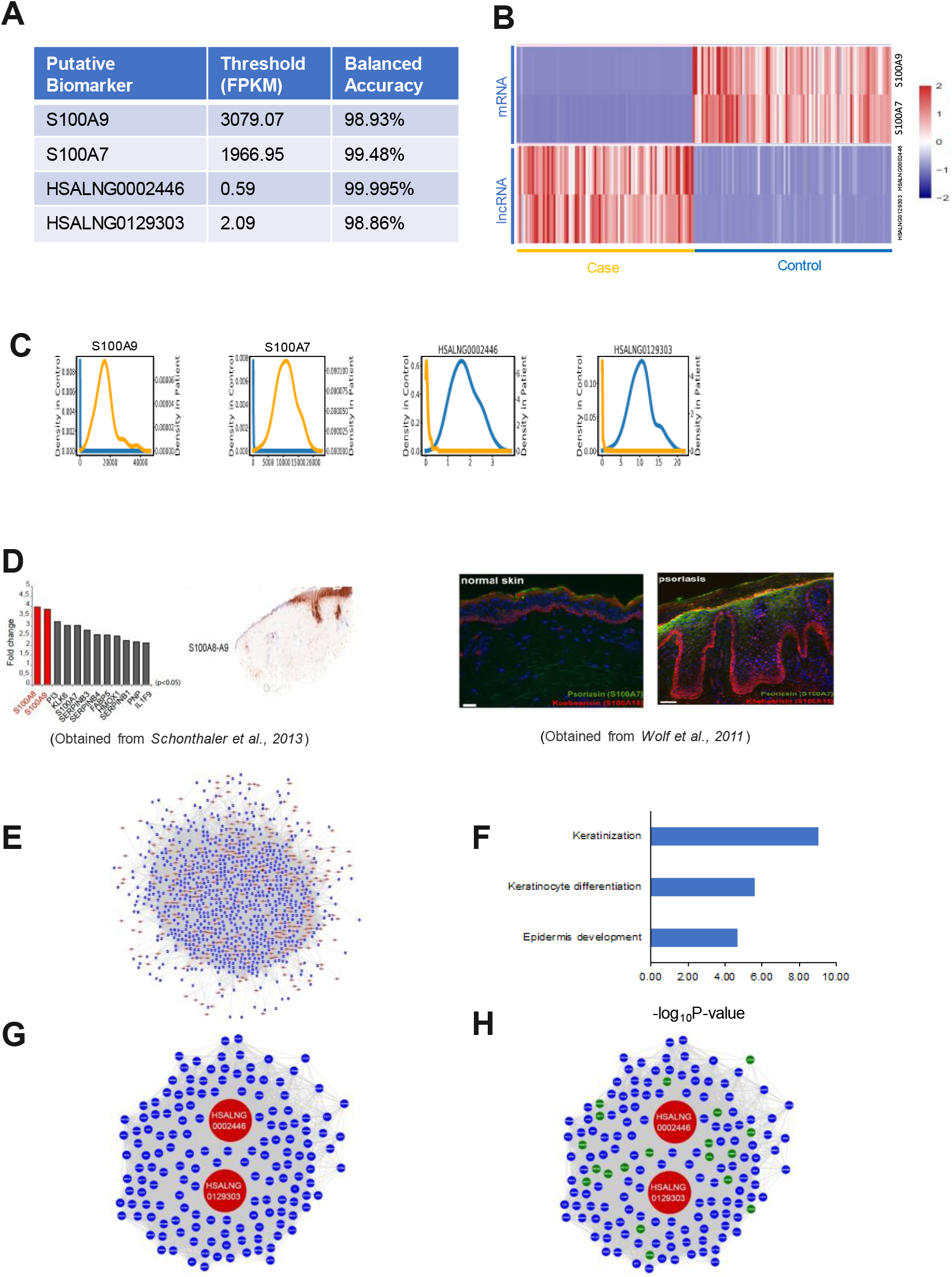
BAMBI discover the biological informative singular mRNA or lncRNA gene as the putative biomarkers for psoriasis diagnosis. We applied BAMBI to the RNA-seq data of skin biospecimens from 92 psoriasis patients and 82 healthy controls. BAMBI found that each of two coding genes (S100A9 and S100A7) and two lncRNAs (HSALNG0002446, HSALNG0129303) would be predictive for psoriasis diagnosis, each with over 98% prediction power. (A) The prediction performance for each of the putative biomarker in diagnosing psoriasis. (B) Heatmaps of the expression profiles for mRNA and lncRNA candidate putative biomarkers across controls and psoriasis specimens. (C) Expression distributions of for these four candidate putative biomarkers across controls and psoriasis specimens. (D) It is well known that the protein products of putative mRNAs biomarkers S100A9 and S100A7) are highly enriched in the psoriatic skins compared normal skins (Broome et al., n.d.; D’Amico et al., 2016; Schonthaler et al., 2013; Silva de Melo et al., 2023; Wolf et al., 2011). (E) The two putative lncRNA-biomarkers are contained in one co-expression cluster. (F) The coding genes of this co-expression cluster containing the putative lncRNA-biomarkers are enriched in the Gene Ontology (GO) categories significantly related to keratinization and epidermis. It indicates these two putative lncRNA-biomarkers (HSALNG0002446 and HSALNG0129303) may contribute to the psoriasis through keratinization and epidermis development. (G) The coding genes (in blue and green) from the cluster in (E) connected to the two putative lncRNA-biomarkers by one-edge are displayed along with the two putative lncRNA-biomarkers (in red). (H)The coding genes that are involved in the keratinization and epidermis development were labelled in green and the other coding genes are labelled in blue.

The four discovered RNA biomarkers shed light significant biological significance in psoriasis. The protein products of two mRNA biomarkers (S100A9 and S100A7) have been shown to be enriched in the skin sections of psoriasis patient vs normal control (**Fig 3C**, (Broome et al., n.d.; D’Amico et al., 2016; chonthaler et al., 2013; il a de Melo et al., 2023; Wolf et al., 2011)). Co-expression network analysis and Gene Ontology enrichment analysis (**Fig 3D-G**) show that the two lncRNA biomarkers are co-expressed with the coding genes involving in the keratinization and epidermis development, which are the key biological process in psoriasis (Iizuka et al., 2004).

### BAMBI identifies putative prognostic biomarkers for acute myeloid leukemia, indictive of patient survival rate

We extended the application of BAMBI in discovering prognostic biomarkers of diseases. To demonstrate this, we applied the BAMBI method to analyze the gene expression profiles of onset conditions of patients with AML, which also have the follow up treatment outcome (**Fig 4A**). Then, we examined if the putative prognostic biomarkers discovered by BAMBI are relevant to the survival rate of the AML patients using an independent evaluation cohort.

**Fig 4:**
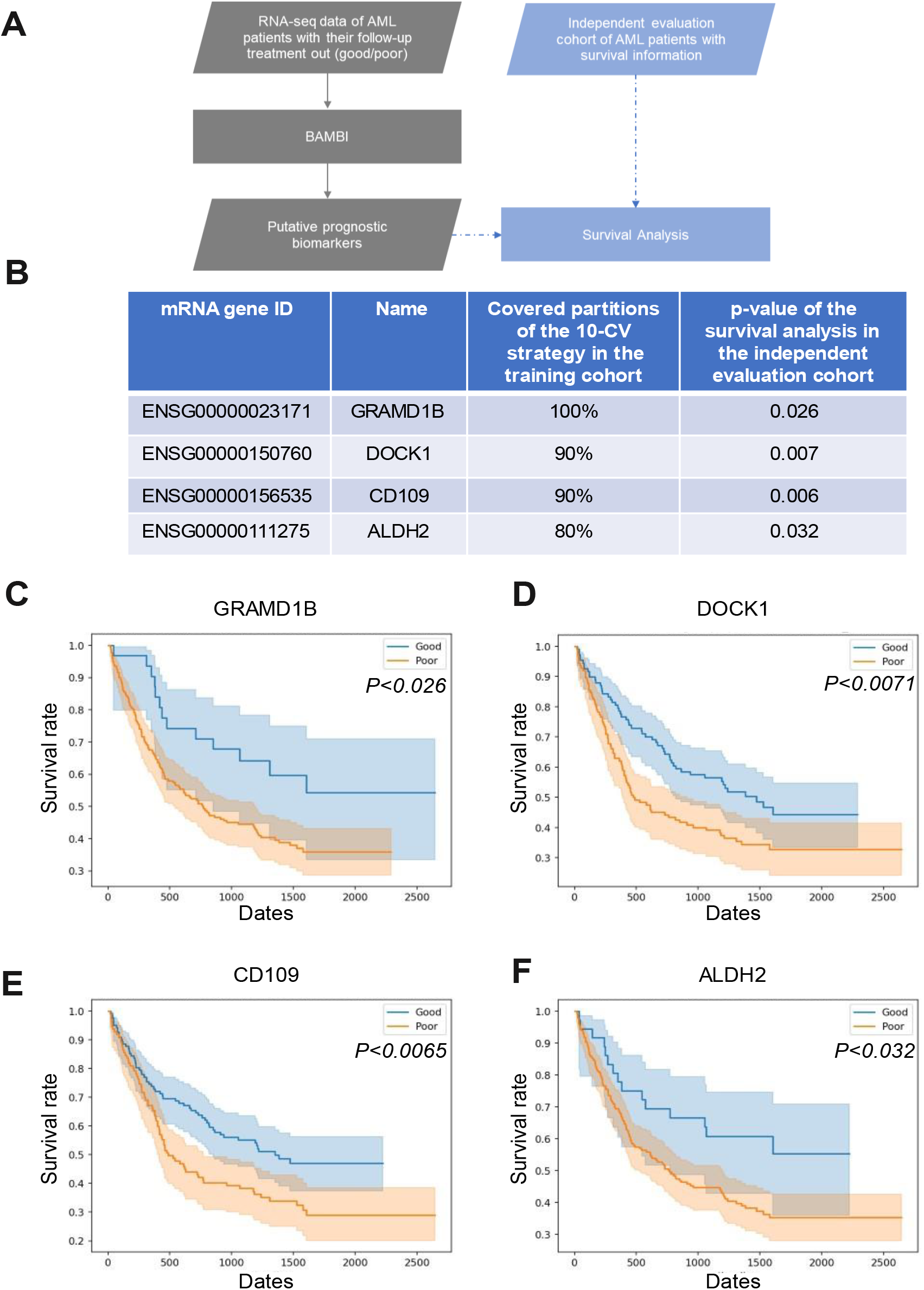
Demonstrating BAMBI in discovering putative RNA prognostic biomarkers for acute myeloid leukemia (AML) (A) The flowchart of applying BAMBI in identifying of RNA prognostic biomarkers for AML and evaluating them in an independent cohort with survival information. (B) The detailed information for each of the putative prognostic biomarker AML. (C-F) The survival curve of overall survival of AML patients with the singular putative prognostic biomarker (panel C: GRAMD1B, panel D: DOCK1, panel E: CD109, panel F: ALDH2) in the evaluation data set. The expression of each of these four putative prognostic biomarkers is significantly indictive of the overall survival of AML patients from the independent evaluation cohort (p<0.05). The p-values were calculated with the logrank test.

Because both training cohort and the independent evaluation cohort did not provide the raw RNA-seq data, instead, provided the mRNA gene expression profiles, it resulted in that lncRNAs were not measured in both cohort and some genes were measured with expression levels in only one of the cohorts. To make this evaluation process consistent, in this application, we used the coding genes with expression profiles in both the training and independent evaluation cohort. Finally, BAMBI discovered nine putative biomarkers which could be identified by at least 8 out of 10 partitions in the training cohort when using 10-CV strategies. Among these nine putative biomarkers, four biomarkers were found to be significantly indictive of the survival rate of AML patients from the independent evaluation cohort (p<0.05) (**Fig 4B-E**). The other five biomarkers did not show significantly relevant to the survival rate of AML patients from the independent cohort; it may be caused by the treatment outcome (good vs poor) measurement in the training cohort does not only rely on patients’ sur i al rate, but also consider other factors, such as age, relapse rate, event-free survival. Younger patients generally have a better overall survival than older patients.

### BAMBI outperforms other methods on both RNA-seq data and microarray data

Several methods have been proposed for RNA biomarker identification. We compared BAMBI with the other state-of-the-art methods coupled with computational tools: BioDiscML (Leclercq et al., 2019), ILRC (Yu et al., 2021), ECMarker (Jin et al., 2021). Because all the other methods cannot process RNA-seq data directly and cannot normalize microarray data by default, we used the RNA gene expression table generated by Phase II of the BAMBI as the input files for the other methods.

We tested the four methods (BAMBI, BioDiscML, ILRC and ECMarker) in both microarray datasets and RNA-seq datasets (**Fig 5A**). Two microarray datasets of colon cancer and prostate cancer were selected in the tool comparison, because they had been previously used in testing the biomarker detection methods. We continued using the two same RNA-seq datasets, that we have used in demonstrating the ability of BAMBI of discovery of singular biomarker. The two RNA-seq datasets include the breast cancer dataset from the TCGA database and the psoriasis dataset from GEO database. For each RNA-seq dataset, we applied the tools to identify mRNA or lncRNA biomarkers, respectively, using the lncBook (Ma et al., 2019) gene annotation information.

**Fig 5:**
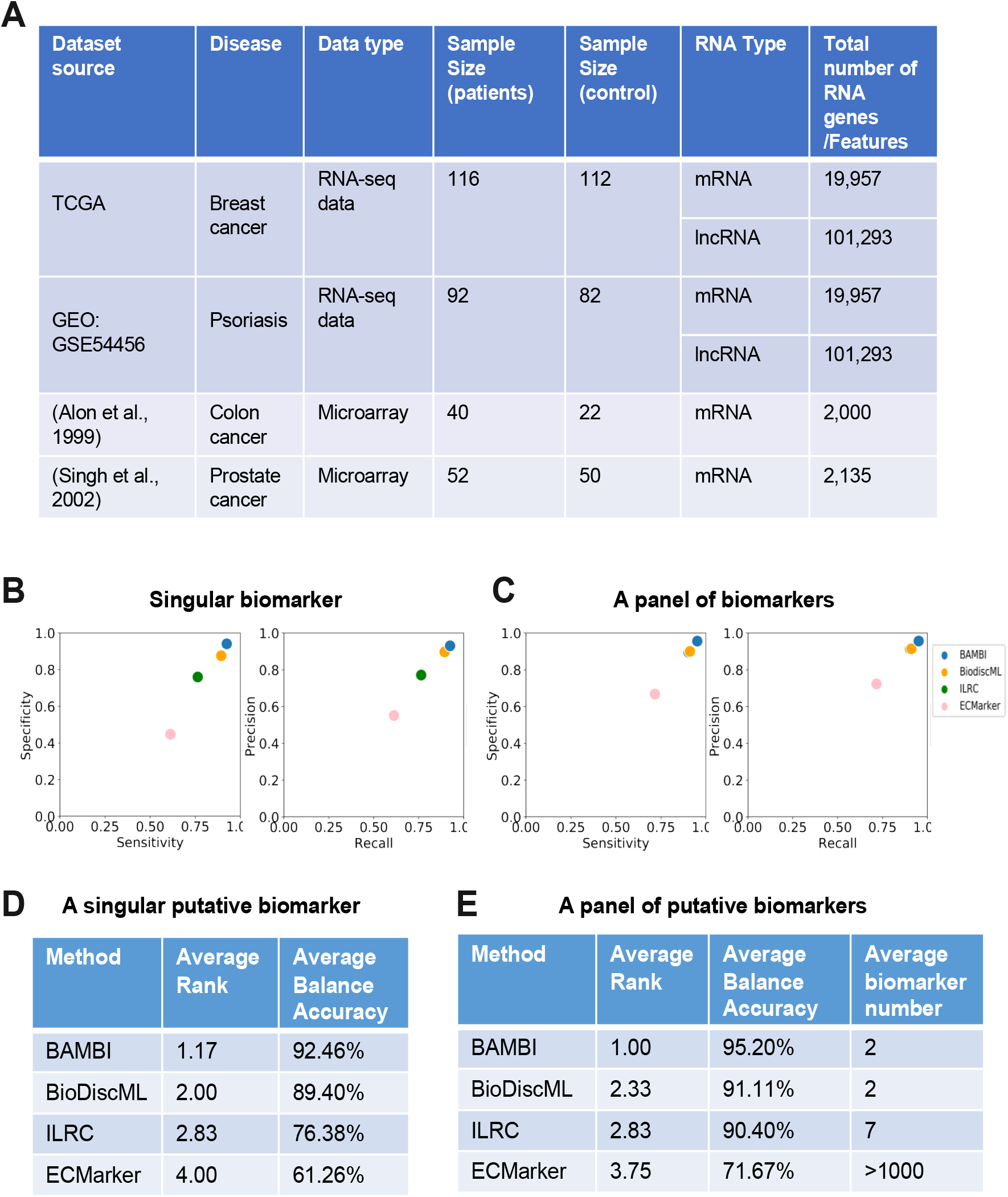
BAMBI outperforms other methods on both RNA-seq data and microarray data. (A)BAMI compared with other methods in two RNA-seq datasets and two microarray datasets. Each RNA-seq dataset were applied to identify mRNA or lncRNA biomarkers (two different cases). (B) The scatter plots show the relationship between performance for each marker gene identified by different methods. The left plot represents the average performance of sensitivity vs specificity, and right plot represents the average performance of precision vs recall for each method across four datasets in sex scenarios (Fig 5A). (C) The scatter plots show the relationship between performance for panel biomarkers identified by different methods. The left plot represents the average performance of sensitivity vs specificity, and right plot represents the average performance of precision vs recall for each method across four datasets in six scenarios (Fig 5A). (D) The average rank of the performance among the four methods (BAMBI, BioDiscML, ILRC and ECMarker) when discovering a singular putative biomarker (E) The average rank of the performance among the four methods (BAMBI, BioDiscML, ILRC and ECMarker) and gene number when applying to identify panel putative biomarkers.

We compared the four methods in detecting two different types of biomarkers: a singular biomarker and a panel of multiple RNAs biomarkers. The output of BAMBI can directly provide these two types of biomarkers, while the other methods only output the best model and related panel biomarker. To evaluate the ability of discovering singular biomarker, here we chose the most important feature of the output best model for the other three method as their singular biomarker.

In our comparative analysis of BAMBI’s performance against other methods, we utilized the cross-validation comparison approach, a crucial consideration given the typically small size of RNA-seq datasets. When splitting such datasets into training and test subsets, the already limited size of the test dataset becomes further reduced. This small size of the test set can lead to the disproportionate influence of outliers or noise on the final evaluation scores. Cross-validation helps mitigate this issue by ensuring a more robust and representative evaluation across the dataset.

Moreover, during the cross-validation process, both feature selection and model training are strictly confined to the training set. This approach guarantees that the model is not exposed to any test set data before the final evaluation phase, thereby avoiding the potential issue of ‘model leakage’ where a model inadvertently gains information from the test data. This careful separation ensures the integrity and generalizability of our evaluation results.

We assessed the performance regarding the specificity vs sensitivity, and precision vs recall (**Fig 5B-C**). The BAMBI method outperformed other methods in both specificity vs sensitivity, and precision vs recall for discovered singular biomarker (**Fig 5B**). Similarly, BAMBI also outperformed other methods regarding the identified panel biomarkers (**Fig 5C**). Overall, BAMBI has the highest balanced accuracy for both singular RNA biomarkers and a panel of multiple RNAs biomarkers (**Fig 5D-E**). While BAMBI achieves high performance, its identified panel biomarkers can still compose of a minimum number of RNA biomarkers. Biomarkers with the minimal gene count as a panel test will facilitate the clinical practice regarding the cost and material availability, and reliability.

## DISCUSSION

We have developed a comprehensive pipeline and software *BAMBI* which integrate statistical methods and machine-learning models to discover coding and noncoding RNA genes as biomarkers from RNA-seq data or microarray data. BAMBI can explore both singular biomarker and panel biomarker. Both singular biomarkers and panel biomarkers discovered by BAMBI can achieve a high prediction power in diagnosing cancer or non-cancerous diseases and outperform other methods. The discovered singular biomarker mode facilitates the easy implementation of biomarkers in clinical practice. These discovered singular biomarkers also significantly shed light on the disease progression as evidence by previous studies for mRNA biomarkers or our co-expression network analysis for lncRNA biomarkers. Our algorithm delivers multiple innovations to solve the limitations and challenges of RNA biomarker discovery.

First, BAMBI overcomes the problem of unreliable and poor predictive performance on independent datasets. To improve the reliability and predictive performance on independent dataset for the discover biomarkers, BAMBI firstly used differentially expressed analysis and overlap area size of distributions across different type of cohorts to significantly reduced the number of RNA gene features, Furthermore, BAMBI imbedded 10-fold based cross-validation strategy to do the feature selection while other methods use the entire dataset to do the feature selection.

Second, BAMBI enhances feature selection by integrating statistical and machine-learning methods. Because one coding or non-coding RNA gene is considered as one feature or one dimension, biomarker discovery process will involve in analyzing hundreds of thousands of coding or non-coding RNA genes. To deal with the extremely high dimension of features when the test datasets are relatively small (<1000 biospecimens) for most transcriptomics datasets, BAMBI integrates statistical and machine-learning methods to select gene expression features.

Third, BAMBI can be adapted to discover prognostic biomarkers as demonstrated on the AML. BAMBI does not require the overall survival information to infer the prognostic biomarker, instead, BAMBI can be conveniently applied to identify prognostic biomarker from the transcriptomics data of patients in on-site condition, as well as their follow-up treatment outcome. It avoids the biases of the prognostic biomarker only indictive of survival rate, while evaluation of therapeutic results depends on multiple factors, including age, survival, relapse, complexity, remission.

Fourthly, BAMBI does not require large-scale dataset as a training cohort. BAMBI has shown high performance on relatively small datasets with less than 100 biospecimens. Due to difficulties in collecting large-scale clinical biospecimens and expensive cost in generating transcriptomics data for large-scale biospecimens, BAMBI become useful in applying to any pilot project of limited resources in biospecimens collection and data generation. Also, BAMBI could be widely used to existing RNA-seq data and microarray dataset of small cohorts.

Fifthly, BAMBI is highly modularized and extensible to directly process RNA-seq transcriptomics data and microarray data. Other methods require users to pre-process RNA-seq data or microarray data to obtain optimal results, whereas BAMBI built in the data preprocessing phase for RNA quantification and normalization directly from RNA-seq or microarray data. It greatly facilitates the easy usage of BAMBI tool for researchers without relevant expertise.

Finally, BAMBI is scalable to discover other non-coding RNA biomarkers, such as circular RNA and micro RNAs, if the RNA expression data includes the expression information of these non-coding RNAs. BAMBI is also able to identify RNA biomarkers while combing all types of coding and non-coding RNA together. While other methods only discovered the most highly expressed gene as biomarker, BAMBI discovered the RNA genes which are most significantly distinguished between cohort independent of their expression level, since BAMBI scales all types of coding and non-coding RNA genes into the same range.

We also implemented BAMBI in parallel computing mode to speed up the data analysis and biomarker discovery. The BAMBI software was implemented in python and wrapped in Docker. We have released the first version of BAMBI tool into the GitHub platform: https://github.com/CZhouLab/BAMBI, along with a user manual.

Together, BAMBI represents a notable advance to identify coding and noncoding RNA biomarkers for diagnosis and prognosis. The discovered RNA biomarkers can accurately diagnose disease or predict the treatment outcome. BAMBI can discover both singular biomarkers and panel biomarkers with minimum gene while maintaining high predictive accuracy, enhancing the biomarkers’ clinical practice. BAMBI tool is applicable to numerous diseases at a transcriptomic-wide scale.

## SOFTWARE AVAILABILITY

The BAMBI software is available on GitHub: https://github.com/CZhouLab/BAMBI with a tutorial.

## Abbreviations

ncRNA: non-coding RNA
lncRNA: long non-coding RNA
mRNA: messenger RNA
GEO: Gene Expression Omics
ML: machine learning
LR: logistic regression
DT: decision tree
KNN: K-nearest neighbors
SVM: support vector machine
AUROC: Area Under the Receiver Operating characteristic Curve
AML: Acute Myeloid Leukemia.

## ACKNOWLEDGEMENTS

Partial manuscript was polished by the AI language model - ChatGPT-4 (July, 2023 version).

## Authors’ contributions

The project was conceived and directed by CZ. The method development and data analysis were performed by PZ with assistance from ZL, FL, SY, JL, and CZ. The results were interpreted by PZ, TH, SV, and CZ. The manuscript was written by PZ and CZ with input from ZL, FL, TH, SY, and SV.

## REFERENCES

Alon, U., Barkai, N., Notterman, D. A., Gish, K., Ybarra, S., Mack, D., & Levine, A. J. (1999). Broad patterns of gene expression revealed by clustering analysis of tumor and normal colon tissues probed by oligonucleotide arrays. In Cell Biology (Vol. 96). http://www.pnas.org.

Anders, S., Pyl, P. T., & Huber, W. (2015). HTSeq-A Python framework to work with highthroughput sequencing data. Bioinformatics, 31(2), 166–169. 10.1093/bioinformatics/btu638

Beylerli, O., Gareev, I., Sufianov, A., Ilyasova, T., & Guang, Y. (2022). Long noncoding RNAs as promising biomarkers in cancer: long non-coding RNAs and cancer. In Non-coding RNA Research (Vol. 7, Issue 2, pp. 66–70). KeAi Communications Co. 10.1016/j.ncrna.2022.02.004

Boieri, M., Malishkevich, A., Guennoun, R., Marchese, E., Kroon, S., Trerice, K. E., Awad, M., Park, J. H., Iyer, S., Kreuzer, J., Haas, W., Rivera, M. N., & Demehri, S. (2022). CD4+ T helper 2 cells suppress breast cancer by inducing terminal differentiation. Journal of Experimental Medicine, 219(7). 10.1084/jem.20201963

Broome, A.-M., Ryan, D., & Eckert, R. L. (n.d.). S100 Protein Subcellular Localization During Epidermal Differentiation and Psoriasis. In Departments of Physiology and Biophysics.

Byron, S. A., Van Keuren-Jensen, K. R., Engelthaler, D. M., Carpten, J. D., & Craig, D. W. (2016). Translating RNA sequencing into clinical diagnostics: Opportunities and challenges. In Nature Reviews Genetics (Vol. 17, Issue 5, pp. 257–271). Nature Publishing Group. 10.1038/nrg.2016.10

D’Amico, F., Skarmoutsou, E., Granata, M., Trovato, C., Rossi, G. A., & Mazzarino, M. C. (2016). S100A7: A rAMPing up AMP molecule in psoriasis. In Cytokine and Growth Factor Reviews (Vol. 32, pp. 97–104). Elsevier Ltd. 10.1016/j.cytogfr.2016.01.002

Demehri, S., Cunningham, T. J., Manivasagam, S., Ngo, K. H., Tuchayi, S. M., Reddy, R., Meyers, M. A., DeNardo, D. G., & Yokoyama, W. M. (2016a). Thymic stromal lymphopoietin blocks early stages of breast carcinogenesis. Journal of Clinical Investigation, 126(4), 1458–1470. 10.1172/JCI83724

Demehri, S., Cunningham, T. J., Manivasagam, S., Ngo, K. H., Tuchayi, S. M., Reddy, R., Meyers, M. A., DeNardo, D. G., & Yokoyama, W. M. (2016b). Thymic stromal lymphopoietin blocks early stages of breast carcinogenesis. Journal of Clinical Investigation, 126(4), 1458–1470. 10.1172/JCI83724

Faratian, D., Sims, A. H., Mullen, P., Kay, C., Um, I. H., Langdon, S. P., & Harrison, D. J. (2011). Sprouty 2 is an independent prognostic factor in breast cancer and may be useful in stratifying patients for trastuzumab therapy. PLoS ONE, 6(8). 10.1371/journal.pone.0023772

FDA. (2016). BEST (Biomarkers, EndpointS, & other Tools) Resource. In Annals of Epidemiology (Vol. 21, Issue 9, pp. 673–687). 10.1016/j.annepidem.2011.02.001

Guennoun, R., Hojanazarova, J., Trerice, K. E., Azin, M., McGoldrick, M. T., Schiferle, E. B., Stover, M. P., & Demehri, S. (2022). Thymic Stromal Lymphopoietin Induction Suppresses Lung Cancer Development. Cancers, 14(9). 10.3390/cancers14092173

Hanafusa, H., Torii, S., Yasunaga, T., & Nishida, E. (2002). Sprouty1 and Sprouty2 provide a control mechanism for the Ras/MAPK signalling pathway. Nature Cell Biology, 4(11), 850–858. 10.1038/ncb867

Hasan, M. M., Alam, M. A., Shoombuatong, W., Deng, H. W., Manavalan, B., & Kurata, H. (2021). NeuroPred-FRL: An interpretable prediction model for identifying neuropeptide using feature representation learning. Briefings in Bioinformatics, 22(6). 10.1093/bib/bbab167

Hasan, M. M., Basith, S., Khatun, M. S., Lee, G., Manavalan, B., & Kurata, H. (2021). Meta-i6mA: An interspecies predictor for identifying DNA N6-methyladenine sites of plant genomes by exploiting informative features in an integrative machine-learning framework. Briefings in Bioinformatics, 22(3). 10.1093/bib/bbaa202

Hasan, M. M., Schaduangrat, N., Basith, S., Lee, G., Shoombuatong, W., & Manavalan, B. (2020). HLPpred-Fuse: Improved and robust prediction of hemolytic peptide and its activity by fusing multiple feature representation. Bioinformatics, 36(11), 3350–3356. 10.1093/bioinformatics/btaa160

Huang, D. W., Sherman, B. T., & Lempicki, R. A. (2009). Systematic and integrative analysis of large gene lists using DAVID bioinformatics resources. Nature Protocols, 4(1), 44–57. 10.1038/nprot.2008.211

Iizuka, H., Takahashi, H., Honma, M., & Ishida-Yamamoto, A. (2004). Unique Keratinization Process in Psoriasis: Late Differentiation Markers Are Abolished because of the Premature Cell Death. In Journal of Dermatology (Vol. 31, Issue 4, pp. 271–276). Japanese Dermatological Association. 10.1111/j.1346-8138.2004.tb00672.x

Jiang, N., Pan, J., Fang, S., Zhou, C., Han, Y., Chen, J., Meng, X., Jin, X., & Gong, Z. (2019). Liquid biopsy: Circulating exosomal long noncoding RNAs in cancer. In Clinica Chimica Acta (Vol. 495, pp. 331–337). Elsevier B.V. 10.1016/j.cca.2019.04.082

Jiao, X., Sherman, B. T., Huang, D. W., Stephens, R., Baseler, M. W., Lane, H. C., & Lempicki, R. A. (2012). DAVID-WS: A stateful web service to facilitate gene/protein list analysis. Bioinformatics, 28(13), 1805–1806. 10.1093/bioinformatics/bts251

Jin, T., Nguyen, N. D., Talos, F., & Wang, D. (2021). ECMarker: Interpretable machine learning model identifies gene expression biomarkers predicting clinical outcomes and reveals molecular mechanisms of human disease in early stages. Bioinformatics, 37(8), 1115–1124. 10.1093/bioinformatics/btaa935

Karimi, B., Dehghani Firoozabadi, A., Peymani, M., & Ghaedi, K. (2022). Circulating long noncoding RNAs as novel bio-tools: Focus on autoimmune diseases. In Human Immunology (Vol. 83, Issues 8–9, pp. 618–627). Elsevier Inc. 10.1016/j.humimm.2022.06.001

Kawazoe, T., & Taniguchi, K. (2019). The Sprouty/Spred family as tumor suppressors: Coming of age. In Cancer Science (Vol. 110, Issue 5, pp. 1525–1535). Blackwell Publishing Ltd. 10.1111/cas.13999

Kim, D., Paggi, J. M., Park, C., Bennett, C., & Salzberg, S. L. (2019). Graph-based genome alignment and genotyping with HISAT2 and HISAT-genotype. Nature Biotechnology, 37(8), 907–915. 10.1038/s41587-019-0201-4

Leclercq, M., Vittrant, B., Martin-Magniette, M. L., Scott Boyer, M. P., Perin, O., Bergeron, A., Fradet, Y., & Droit, A. (2019). Large-scale automatic feature selection for biomarker discovery in high-dimensional omics data. Frontiers in Genetics, 10(MAY). 10.3389/fgene.2019.00452

Li, Y., Ge, X., Peng, F., Li, W., & Li, J. J. (2022). Exaggerated false positives by popular differential expression methods when analyzing human population samples. Genome Biology, 23(1). 10.1186/s13059-022-02648-4

Love, M. I., Huber, W., & Anders, S. (2014). Moderated estimation of fold change and dispersion for RNA-seq data with DESeq2. Genome Biology, 15(12). 10.1186/s13059-014-0550-8

Lundberg, S. M., Allen, P. G., & Lee, S.-I. (2017). A Unified Approach to Interpreting Model Predictions. https://github.com/slundberg/shap

Ma, L., Cao, J., Liu, L., Du, Q., Li, Z., Zou, D., Bajic, V. B., & Zhang, Z. (2019). Lncbook: A curated knowledgebase of human long non-coding rnas. Nucleic Acids Research, 47(D1), D128–D134. 10.1093/nar/gky960

Ratti, M., Lampis, A., Ghidini, M., Salati, M., Mirchev, M. B., Valeri, N., & Hahne, J. C. (2020). MicroRNAs (miRNAs) and Long Non-Coding RNAs (lncRNAs) as New Tools for Cancer Therapy: First Steps from Bench to Bedside. In Targeted Oncology (Vol. 15, Issue 3, pp. 261–278). Adis. 10.1007/s11523-020-00717-x

Rifai, N., Gillette, M. A., & Carr, S. A. (2006). Protein biomarker discovery and validation: The long and uncertain path to clinical utility. In Nature Biotechnology (Vol. 24, Issue 8, pp. 971–983). 10.1038/nbt1235

Ritchie, M. E., Phipson, B., Wu, D., Hu, Y., Law, C. W., Shi, W., & Smyth, G. K. (2015). Limma powers differential expression analyses for RNA-sequencing and microarray studies. Nucleic Acids Research, 43(7), e47. 10.1093/nar/gkv007

Robinson, M. D., McCarthy, D. J., & Smyth, G. K. (2009). edgeR: A Bioconductor package for differential expression analysis of digital gene expression data. Bioinformatics, 26(1), 139–140. 10.1093/bioinformatics/btp616

Schonthaler, H. B., Guinea-Viniegra, J., Wculek, S. K., Ruppen, I., Ximénez-Embún, P., Guío-Carrión, A., Navarro, R., Hogg, N., Ashman, K., & Wagner, E. F. (2013). S100A8-S100A9 Protein Complex Mediates Psoriasis by Regulating the Expression of Complement Factor C3. Immunity, 39(6), 1171–1181. 10.1016/j.immuni.2013.11.011

Silva de Melo, B. M., Veras, F. P., Zwicky, P., Lima, D., Ingelfinger, F., Martins, T. V., da Silva Prado, D., Schärli, S., Publio, G., Hiroki, C. H., Melo, P. H., Saraiva, A., Norbiato, T., Lima, L., Ryffel, B., Vogl, T., Roth, J., Waisman, A., Nakaya, H. I., … Alves-Filho, J. C. (2023). S100A9 Drives the Chronification of Psoriasiform Inflammation by Inducing IL-23/Type 3 Immunity. Journal of Investigative Dermatology, 143(9), 1678–1688.e8. 10.1016/j.jid.2023.02.026

Singh, D., Febbo, P. G., Ross, K., Jackson, D. G., Manola, J., Ladd, C., Tamayo, P., Renshaw, A. A., D’amico, A. V, Richie, J. P., Lander, E. S., Loda, M., Kantoff, P. W., Golub, T. R., & Sellers, W. R. (2002). Gene expression correlates of clinical prostate cancer behavior. http://www-genome.wi.mit.edu/MPR/prostate

Tischler, G., & Leonard, S. (2014). biobambam: tools for read pair collation based algorithms on BAM files. http://www.scfbm.org/content/9/1/13

Walker, J. M. (2011). METHODS IN MOLECULAR BIOLOGY. http://www.springer.com/series/7651

Wang, Y., Zhang, M., & Zhou, F. (2020). Biological functions and clinical applications of exosomal long non-coding RNAs in cancer. In Journal of Cellular and Molecular Medicine (Vol. 24, Issue 20, pp. 11656–11666). Blackwell Publishing Inc. 10.1111/jcmm.15873

Weinstein, J. N., Collisson, E. A., Mills, G. B., Shaw, K. R. M., Ozenberger, B. A., Ellrott, K., Sander, C., Stuart, J. M., Chang, K., Creighton, C. J., Davis, C., Donehower, L., Drummond, J., Wheeler, D., Ally, A., Balasundaram, M., Birol, I., Butterfield, Y. S. N., Chu, A., … Kling, T. (2013). The cancer genome atlas pan-cancer analysis project. Nature Genetics, 45(10), 1113–1120. 10.1038/ng.2764

Wolf, R., Ruzicka, T., & Yuspa, S. H. (2011). Novel S100A7 (psoriasin)/S100A15 (koebnerisin) subfamily: Highly homologous but distinct in regulation and function. In Amino Acids (Vol. 41, Issue 4, pp. 789–796). 10.1007/s00726-010-0666-4

Xi, X., Li, T., Huang, Y., Sun, J., Zhu, Y., Yang, Y., & Lu, Z. J. (2017a). RNA biomarkers: Frontier of precision medicine for cancer. In Non-coding RNA (Vol. 3, Issue 1). MDPI AG. 10.3390/ncrna3010009

Xi, X., Li, T., Huang, Y., Sun, J., Zhu, Y., Yang, Y., & Lu, Z. J. (2017b). RNA biomarkers: Frontier of precision medicine for cancer. In Non-coding RNA (Vol. 3, Issue 1). MDPI AG. 10.3390/ncrna3010009

Yu, K., Xie, W., Wang, L., & Li, W. (2021). ILRC: a hybrid biomarker discovery algorithm based on improved L1 regularization and clustering in microarray data. BMC Bioinformatics, 22(1). 10.1186/s12859-021-04443-7

