## Supplement Materials and Method for "BAMBI: Integrative *b*iostatistical and *a*rtificial-intelligence *m*odels discover coding and non-coding RNA genes as *bi*omarkers"

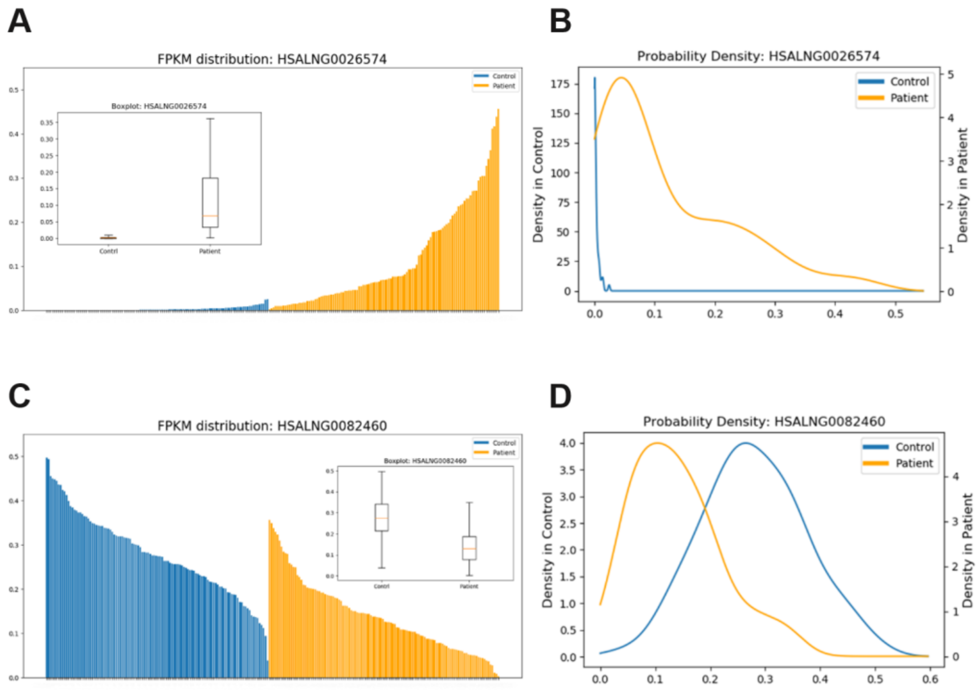


**Supplement Fig 1: High distribution overlap genes filter.** Typically, people use the mean or median difference to evaluate the expression heterogeneity. In BAMBI, we also add the estimate distribution overlap between groups as a criterion. Compared to mean and median differences, estimate distribution overlap can be more comprehensively and accurately describe whether the gene expression is heterogeneous between groups.

(A B) are the expressions of two noncoding RNAs from the TCGA breast cancer dataset. Both the group mean difference and median difference of gene HSALNG0082460 (mean difference: 0.14; median difference: 0.15) are more significant than gene HSALNG0026574 (mean difference: 0.11; median difference: 0.07). However, from the observations of the expression FPKM data distribution and boxplot, in gene HSALNG0082460, a considerable expression overlap exists between the control and patient groups. If we use such kinds of genes as a biomarker to predict disease, there will be an extensive uncertain expression range that we cannot decide it is from a patient or a non-patient. Compared to gene HSALNG0082460, gene HSALNG0026574 has minor between-group mean and median differences. However, its estimated distribution overlap (C) is only 1.7% (compared to the 31.2% of HSALNG0082460 (D)), and its uncertain range is much narrower.

The biomarker gene with low distribution overlap between groups will be more practical. Take the diagnostic biomarker as an example. If the biomarker gene has a clear gap between patient and healthy control groups, the doctor can use this expression gap as a threshold to predict the disease status directly.

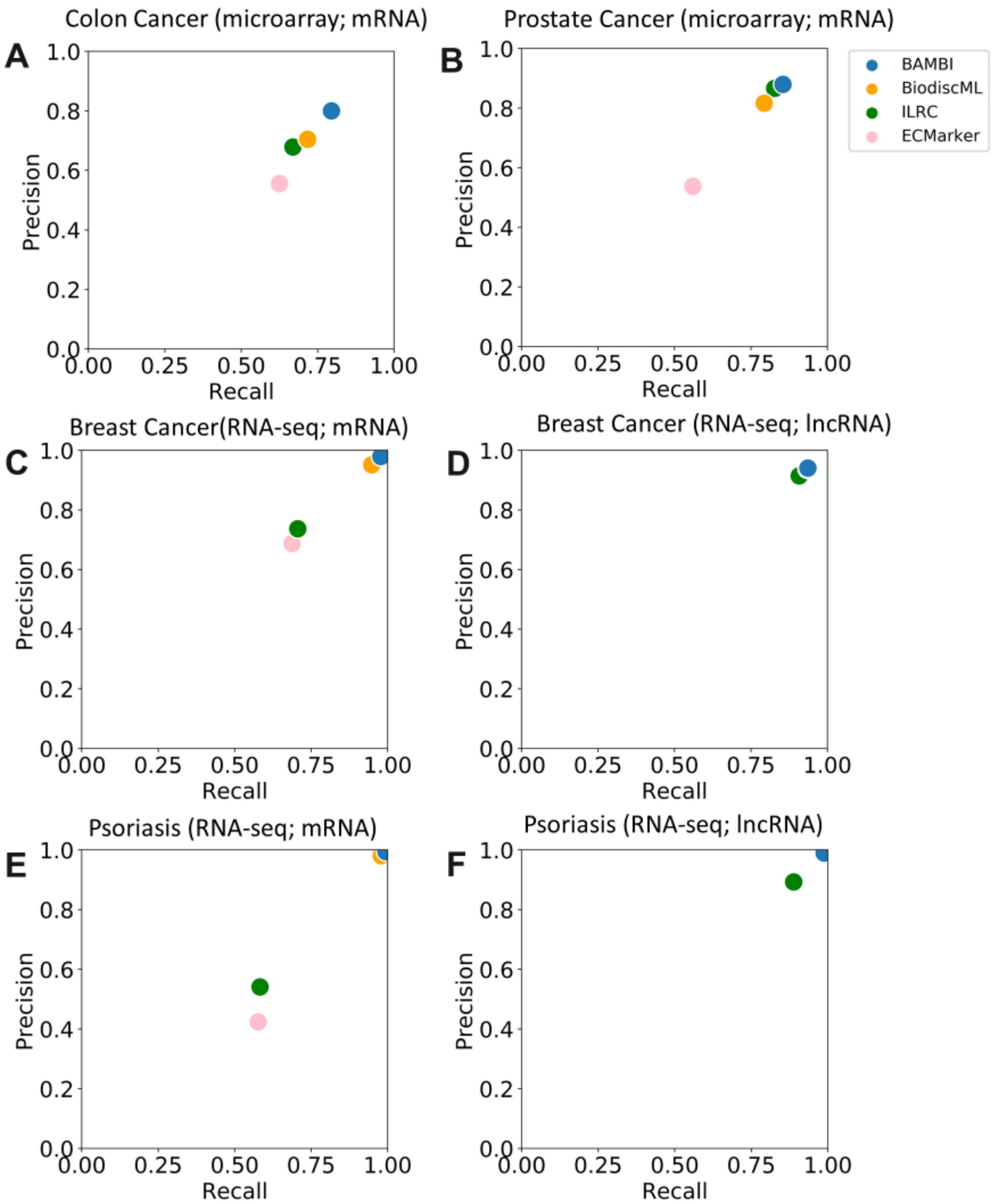


**Supplement Fig 2: BAMBI surpasses competing methods in achieving higher Precision vs Recall rates for identifying singular biomarkers, demonstrating superior performance in both RNA-seq and microarray data analyses.** (A-F) The scatter plots show the relationship between performance for each singular biomarker gene identified by different methods. Each plot represents the performance of precision vs recall for each method in different scenarios.

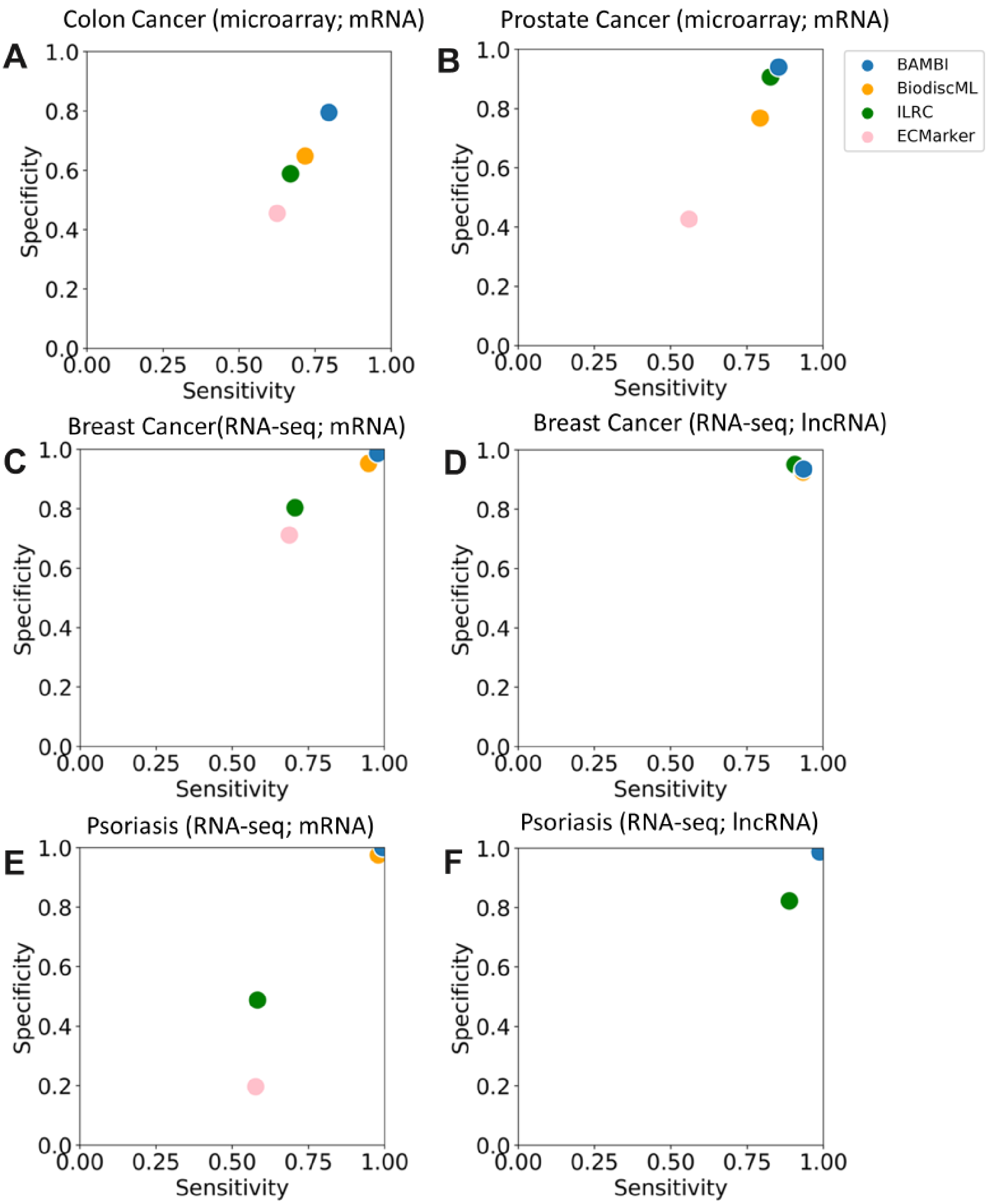


**Supplement Fig 3: BAMBI surpasses competing methods in achieving higher Sensitivity vs Specificity rates for identifying singular biomarkers, demonstrating superior performance in both RNA-seq and microarray data analyses.** (A-F) The scatter plots show the relationship between performance for each singular biomarker gene identified by different methods. Each plot represents the performance of precision vs recall for each method in different scenarios.

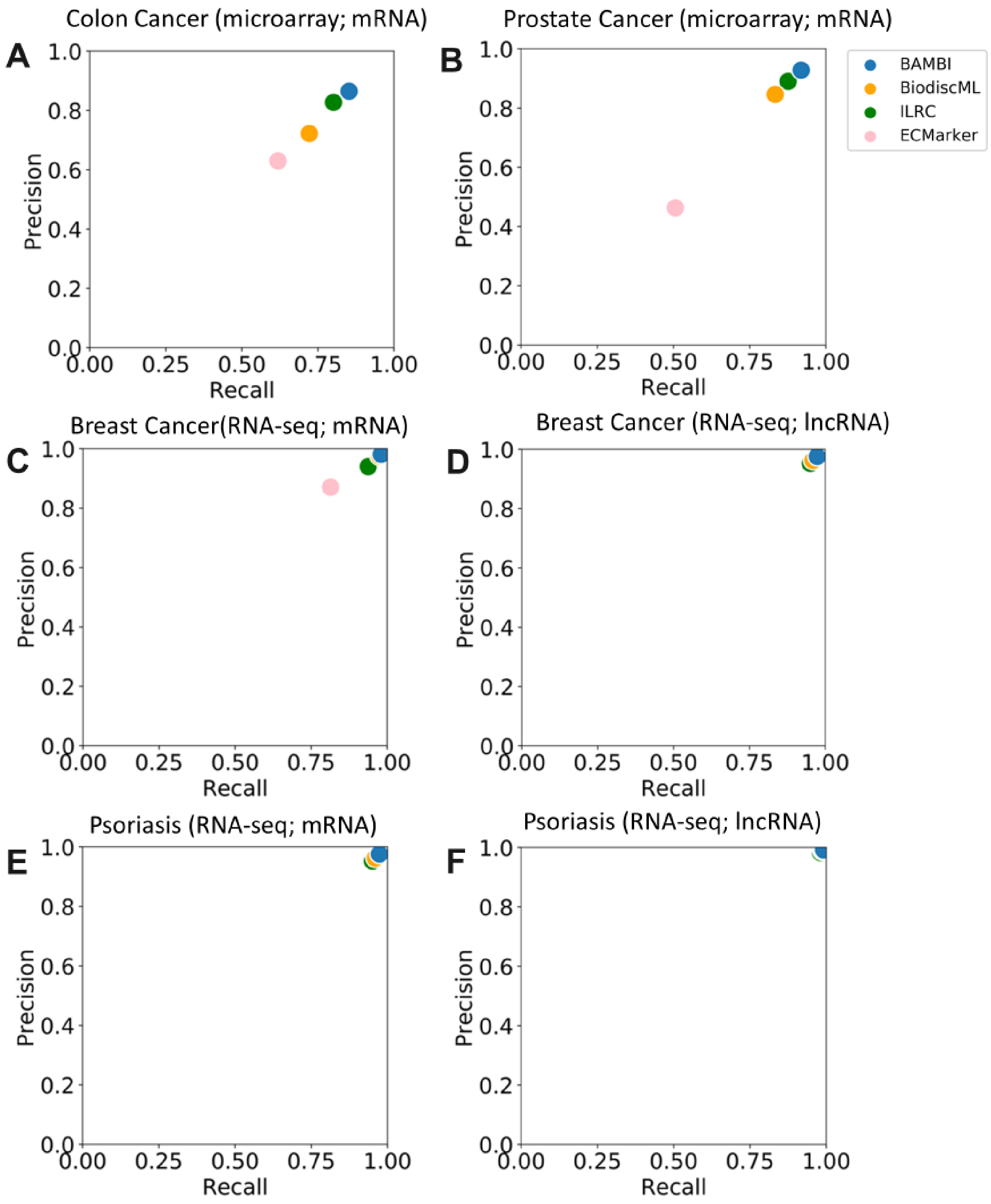


**Supplement Fig 4: BAMBI surpasses competing methods in achieving higher Precision vs Recall rates for identifying panel biomarkers, demonstrating superior performance in both RNA-seq and microarray data analyses.** (A-F) The scatter plots show the relationship between performance for each panel biomarker identified by different methods. Each plot represents the performance of precision vs recall for each method in different scenarios.

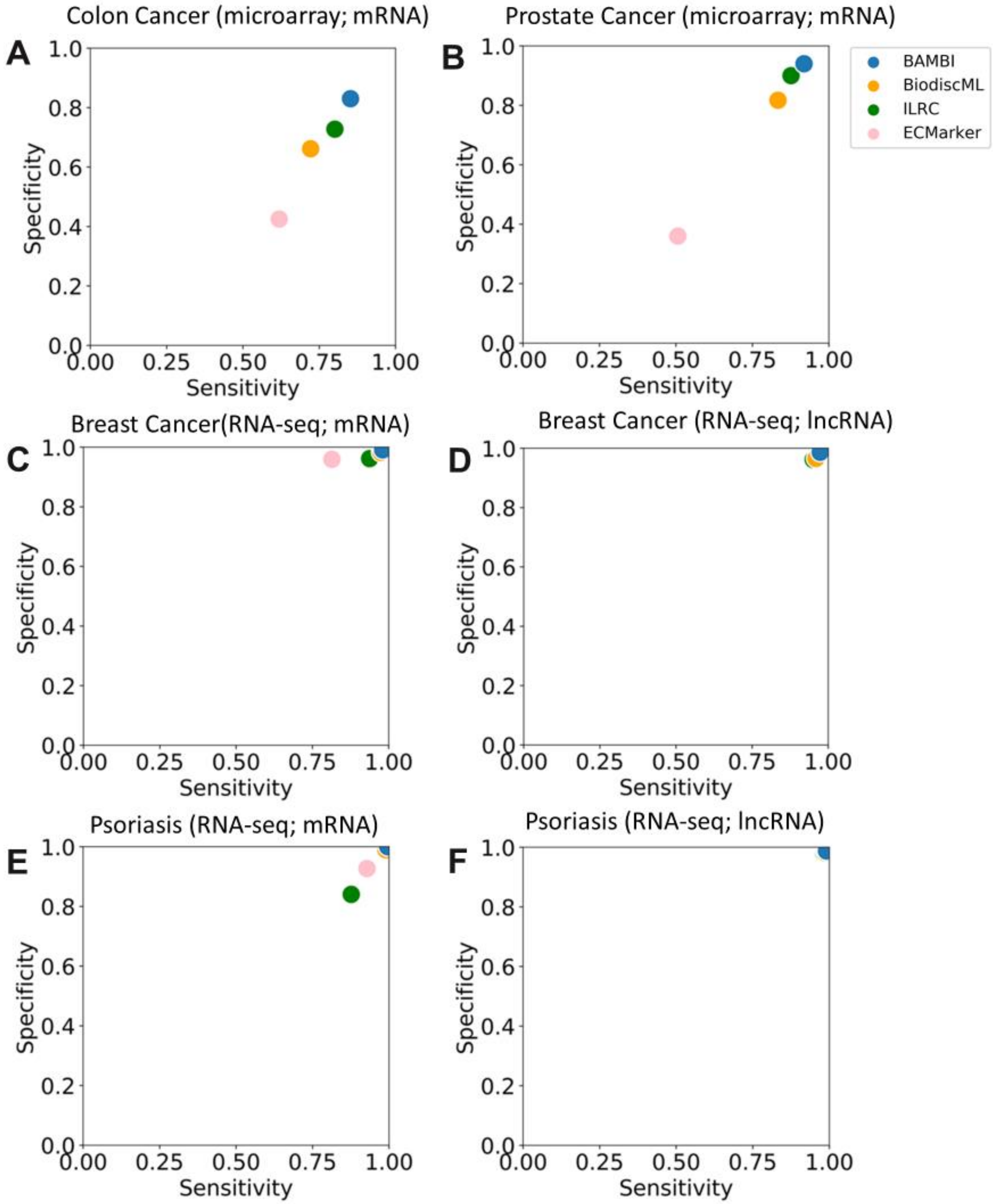


**Supplement Fig 4: BAMBI surpasses competing methods in achieving higher Sensitivity vs Specificity rates for identifying panel biomarkers, demonstrating superior performance in both RNA-seq and microarray data analyses.** (A-F) The scatter plots show the relationship between performance for each panel biomarker identified by different methods. Each plot represents the performance of precision vs recall for each method in different scenarios.
